# SSR42 Enhances the Hemolytic Capacity of *Staphylococcus aureus* by Stabilizing the *hla* mRNA

**DOI:** 10.64898/2026.07.23.740365

**Authors:** Mary-Elizabeth Jobson, Brooke R. Tomlinson, Jessica K. Jackson, Lindsey N. Shaw

**Author notes:** Address correspondence to Lindsey N. Shaw, PhD.

## Abstract

*Staphylococcus aureus* utilizes a complex regulatory network to precisely control a vast array of virulence factors and facilitate infection. One such regulator, the long regulatory RNA SSR42, is emerging as a key modulator of virulence factor abundance and pathogenesis. Previous work has shown that a primary role for SSR42 is controlling hemolytic behavior through the positive regulation of α-toxin (*hla*/Hla). Herein, we confirm that loss of SSR42 limits hemolytic capacity due to reduced Hla production. Others have suggested that this occurs through the SaeRS two-component system; however, using epistasis experiments, we reveal that SSR42 and SaeRS function independently to control Hla activity. Using a two-plasmid system optimized to detect regulatory RNA-mediated control in Gram-positive bacteria, we demonstrate that SSR42 directly enhances Hla production through the *hla* 5’ untranslated region. Using interaction prediction and targeted mutagenesis, we identify a discrete site upstream of the *hla* ribosome-binding site that is required for SSR42 binding and hemolytic activity. Transcriptional arrest experiments further show that SSR42 stabilizes the *hla* mRNA, increasing its half-life and promoting downstream toxin production. Finally, electrophoretic mobility shift assays confirm a specific interaction between SSR42 and the *hla* 5’ UTR. Collectively, these findings establish SSR42 as a major post-transcriptional regulator of *S. aureus* virulence and reveal a novel RNA-mediated mechanism that promotes α-toxin production through stabilization of the *hla* transcript.

## Introduction

In the context of secreted factors, *S. aureus* exhibits three key pillars of virulence: cytolysis, proteolysis, and hemolysis(1, 2). Hemolytic ability is one of the most prototypical behaviors of *S. aureus* and has been primarily shown to function through alpha-toxin, Hla (3). Hla is capable of lysing a variety of host cell types, exerts immunomodulatory effects, and has been shown to be required for infection in animal models of dermonecrosis, pneumonia, sepsis, and more (4-16).

Regulatory control over *hla* expression is carried out by a multitude of elements, including two-component systems (TCSs), alternative sigma factors, DNA binding transcription factors, and regulatory RNAs (often referred to as sRNAs (1, 17−21)). The most well studied contributor to the regulation of hemolysis is the accessory gene regulator (Agr), a TCS and quorum sensing system which functions primarily to control the switch during infection towards a more toxic, invasive phenotype(17, 18). This system also encodes RNAIII, a regulatory RNA-based effector that modulates virulence via direct binding to mRNA transcripts to affect their translation(19). In the context of Hla, RNAIII is capable of impacting its activity both on a local and global scale. On a direct level, RNAIII binds to the *hla* mRNA to activate its translation through liberation of the ribosome binding site (RBS). More globally, RNAIII has been shown to also control *hla* expression via action on the well characterized <u>r</u>epressor <u>o</u>f <u>t</u>oxins (Rot) transcription factor (20, 21). In this scenario, RNAIII binds to the *rot* mRNA RBS, preventing its translation and promoting recruitment of RNase III that degrades the mRNA. This consequently halts production of the *hla-* repressing Rot protein.

While RNAIII is the most well-studied regulatory RNA, not only in *S. aureus* but in bacteria in general, recent work has revealed a multitude of RNAs that play pivotal roles in complex regulatory systems, ranging from antisense regulatory transcripts to *trans* acting RNAs (22, 23). Virulence-related sRNAs in *S. aureus* include the Spr-group, such as SprC and SprD, which regulate the major autolysin *atl* and the adhesive virulence factor *sbi*, respectively (24, 25). Another family of regulatory RNAs is the Teg RNAs, which includes Teg41 that impacts hemolytic ability through the PSMs and plays a role in promoting pathogenesis (26, 27). The regulation of virulence is invariably linked to the metabolic status of the cell, which has been shown to also be tightly controlled by regulatory RNAs. One such example is RsaC, a ∼1,100-nt transcript that coordinates manganese homeostasis while enhancing the oxidative stress response, leading to increased ability to survive confrontation with the host immune system (28).

Previous work by our group characterized a unique RNA-regulator, SSR42, which is the second most abundant transcript in the *S. aureus* cell during stationary phase, second only to RNAIII (29). Using multi-omics technology, we identified a broad and complex SSR42 regulon that encompasses a multitude of virulence factors, including secreted proteases and toxins. We also uncovered a unique pathway in which SSR42 responds to the presence of neutrophils and upregulates *lukAB* expression to facilitate cytotoxicity. First characterized by Morrison et al. from a larger group of Small Stable RNAs (SSRs), SSR42 has been subject to discovery and rediscovery by numerous groups, and is also named Teg27, sRNA 363, sRN_4470, and RSaX28 from various sources (30-36). It has previously been shown that SSR42 exhibits significantly enhanced stability and abundance, particularly during stationary phase, contains no functional ORFs, and influences virulence factor production in strains UAMS-1 and USA300 LAC (35, 36). It has also been observed that the SSR42 nucleotide sequence is almost completely conserved (98%) across strains of *S. aureus*, with only two single nucleotide variants found in the 819-nt long transcript from ST30 isolates (36). Other foundational work showed that SSR42 plays a role in controlling hemolysis and may respond to the presence of antibiotics, however direct targets and/or regulators have yet to be fully elucidated in these works (35). Herein, we show that SSR42 is a novel, direct regulator of hemolytic activity that functions independently of previously characterized regulatory systems. We determine that SSR42 stabilizes the *hla* mRNA at the 5’ UTR, resulting in enhanced downstream translation. We use mutagenesis to probe RNA-RNA engagement between these two molecules, identifying a region of interaction just upstream of the *hla* RBS that facilitates upregulation of hemolytic ability. Taken together, we have identified a novel form of hemolysis regulation that functions independently of known mechanisms, solidifying SSR42 as an important regulator of the virulence process.

## Materials and Methods

### Bacterial strains and growth conditions

Overnight cultures were routinely grown at 37°C with shaking at 250 rpm in 5mL of tryptic soy broth (TSB) for *S. aureus* or lysogeny broth (LB) for *E*.*coli*. When required, media was supplemented with the following antibiotics: *E*.*coli:* 100µg/mL ampicillin; *S. aureus:* 10µg/mL chloramphenicol, 5µg/mL erythromycin, 25µg/mL lincomycin, and 5µg/mL tetracycline. To obtain synchronous growth of *S. aureus*, overnight cultures were diluted 1:100 into fresh TSB and grown for 3h before being standardized to a uniform OD_600_ of 0.05 in fresh TSB. Unless otherwise described, cell pellets were harvested via centrifugation at 4000 × *g* for 10 minutes.

### Strain construction

All bacterial strains used in this study are listed in **Table S1**, plasmids in **Table S2**, and primers in **Table S3**. Deletion of SSR42 as well as construction of complementing strains was described previously (29). Deletion of *sarR* was previously described (37), while the *saeR, rsp*, and *sarZ* mutations were obtained from the Nebraska Transposon Mutant Library (NTML) (38). Phage transductions were performed as previously described (39) to mobilize mutations into the appropriate genetic background, and all strains were confirmed using gene specific primers.

### Hemolysis assays

Hemolysis was measured using two methods. First, by isolating a single colony and inoculating (via an agar stab) TSA containing 5% sheep’s blood. Secondly, liquid bacterial cultures were grown in biological triplicate overnight and the OD_600_ of all samples were standardized to each other. Following this, 7µl of standardized culture was spot plated onto TSA containing 5% sheep’s blood and grown overnight. Clearing of the blood substrate around bacterial growth was quantified using ImageJ.

### Reporter, overexpression, and mutagenesis constructs

Transcriptional reporters were constructed utilizing the pXEN-1 vector which fuses a promoter of interest with the *luxABCDE* operon (40). The SSR42 and *hla* promoters were amplified using OL4526/OL7967 and OL843/844, respectively and cloned into pXen-1 using restriction enzymes BamHI and EcoRI. Translational fusions were created utilizing vectors optimized for monitoring sRNA regulation in Gram-positive bacteria (41). The *hla* UTR and a few residues of the coding region were amplified using OL7424/7481 and cloned into pCN33 using restriction enzymes Bglll and EcoRV. The SSR42 expression vector used in this system was previously described (42). Overexpression of *sarR* was achieved by cloning a promoterless *sarR* behind a cadmium-inducible promoter in the pJB67 vector. *sarR* was amplified using OL6637/6640 and cloned into pJB67 using restriction enzymes Ndel and EcoRI. Mutagenesis of the SSR42 complementation plasmids was performed using the NEB Q5 mutagenesis kit and manufacturer instructions. Mutagenesis of SSR42 was performed using either primers OL8046/8047 or OL8044/8045 for sequence scrambling or deletion, respectively. All clones were transformed into chemically competent *E*.*coli* DH5α before being electroporated into *S. aureus* RN4220 and finally transduced into relevant *S. aureus* strains. All constructs were confirmed via PCR and sequencing.

### RT-qPCR

Real-Time Quantitative Reverse Transcription PCR (RT-qPCR) was performed as previously described (43). In brief, strains were synchronized and standardized to an OD_600_ of 0.05 before being grown for 15h and total RNA isolated from cell pellets. Samples were reverse transcribed using an iScript cDNA synthesis Kit (Biorad) and RT-qPCR was performed using gene-specific primers **(Table S3)** and TB Green Premix Ex Taq (Takara). Expression levels were normalized to 16S rRNA and fold change of expression was determined using the 2−ΔΔCT method.

### Transcriptional Arrest

Transcriptional arrest and determination of RNA half-life was performed as described (43). In brief, bacterial strains were synchronized for 3h before being standardized to an OD_600_ of 0.05 and grown for 15h. After this time, 5mLs of culture was collected, immediately combined with ice-cold PBS, and cells were harvested via refrigerated centrifugation. Rifampin was then added to a final concentration of 250µg/mL to each culture and samples were taken at 5, 10, 15, 30, and 45 minutes posttreatment. RNA isolation and qPCR were performed as described above for each sample, with quantification of RNA abundance at each timepoint calculated using the 2−ΔΔCT relative to the initial RNA abundance at t= 0. A one phase decay curve was generated using Graph pad Prism to determine the decay rate *k*, which was subsequently used to calculate the RNA half-life using the equation t_½_ = ln(2)/k.

### Transcriptional and translational reporter assays

To measure promoter activity, bacterial strains containing luciferase-based promoter fusions were grown overnight in biological triplicate. Following this, cultures were synchronized for 3h before being standardized to an OD_600_ of 0.05 into a black-walled 96-well plate containing TSB and the appropriate selecting antibiotics. Finally, using a Biotek Cytation 5 Plate reader, luminescence and OD_600_ measurements were taken every 15 min for 18h. To measure translational control of SSR42, strains containing the target mRNA-GFP fusion (pCN33) were expressed either in the presence or absence of SSR42 *in trans* (plCS3). Bacterial cultures were prepared as described above, however following standardization, fluorescence (excitation: 485/20, emission: 528/20, Gain: 75) and OD_600_ were measured every 15 min for 18h. Strains were normalized by growth via OD_600_.

### Electrophoretic mobility shift assays (EMSAs)

RNA-RNA EMSAs were performed as previously described (42). In brief, RNA fragments of interest were transcribed *in vitro* using a T7 RNA polymerase promoter and the MAXIscript T7 kit (ThermoFisher). SSR42 was amplified using OL7499/7500 while fragments of the *hla* UTR and *splE* UTR were amplified using primers OL8117/8118 and OL7523/7524, respectively. *in vitro* transcription was performed according to manufacturer instructions using 100ng of template DNA, ^32^P-labeled UTP for the target mRNAs, and unlabeled UTP for SSR42. Following transcription, TurboDNAse was added, and reactions were purified using a Nucleotide Clean-up Kit (Qiagen). RNA-RNA binding reactions were prepared using labeled *hla* and SSR42 at varying concentrations. These were then incubated at 90°C for 2 minutes, followed by incubation at RT for 30 minutes. Native acrylamide gels were equilibrated on ice for 30 minutes in 0.5x TBE before binding reactions were mixed with RNA loading dye and run at 100V until the dye front moved ¾ of the way through the gel. Gels were dried and affixed to filter paper before being visualized via autoradiography.

## Results

### SSR42 regulates hemolytic activity in *S. aureus* independently of the SaeRS TCS

Previous work by others has demonstrated that SSR42 mutants are impaired in their ability to lyse rabbit- (USA300 LAC, ST8) (36) and sheep- (strain 6850, ST50) (35) erythrocytes. This is driven by a role for SSR42 in positively stimulating production of alpha-hemolysin (36). Work performed in the *S. aureus strain* 6850 suggests that this interaction is indirect, and likely occurs through the SaeRS TCS, a strong regulator of Hla activity (35). Previously, we have validated decreased *hla*/Hla abundance in a USA300 SSR42 mutant at the transcript and protein level (29)(42) and confirm that change herein via western blot **(Figure 1A)**. Importantly, however, our published RNAseq dataset (29) shows no alteration in *saeRS* transcription upon SSR42 deletion. To investigate this further, we first quantified *saeR* and *hla* transcript levels in the USA300 HOU WT and SSR42 mutant strains using qPCR. In so doing we noted that *saeR* was not significantly altered in expression in the mutant, whilst the *hla* mRNA decreased ∼3-fold in abundance, confirming previous transcriptomic and proteomic datasets **(Figure 1B)**. We next sought to further confirm that SaeRS has no impact on SSR42-mediated control of Hla at the level of activity. To that end, we created a series of mutant strains and tested their ability to lyse sheep erythrocytes - a specific measure of Hla activity. As expected, an SSR42 mutant exhibits a significant decrease in hemolytic ability, which is fully complemented upon re-introduction of SSR42 on a plasmid **(Figure 1CD)**. When assessing the *saeR* mutant, we also see hemolysis decline compared to the parental strain, again as expected (45−47). Interestingly, when we create a double mutant by deleting SSR42 in the *saeR*-deficient strain, we observe an additive effect, with a significant additional decrease in hemolytic activity noted. Importantly, upon reintroduction of SSR42 (underthe control of its native promoter) on a plasmid into this double mutant, we observed a reversal of this phenotype **(Figure 1CD)**. This demonstrates that SSR42 exerts its own SaeRS independent effect on hemolytic ability via action on Hla. As a final query to uncover if SSR42 is at all connected to SaeRS in a regulatory pathway, we investigated if SSR42’s transcription is under control of SaeRS. To do so, we fused the SSR42 promoter with luciferase and expressed this in an *saeR* mutant strain, comparing it to wild-type. Here we observed no significant change in promoter activity, most notably at the 15h timepoint used for all experiments assessing *hla* abundance and activity **(Figure S1)**. Thus, we posit that SSR42 has its own independent effect on Hla activity and can induce hemolysis in the absence of SaeRS, supporting the notion that SSR42 oversees a novel, direct pathway controlling Hla production.

**Figure 1.**
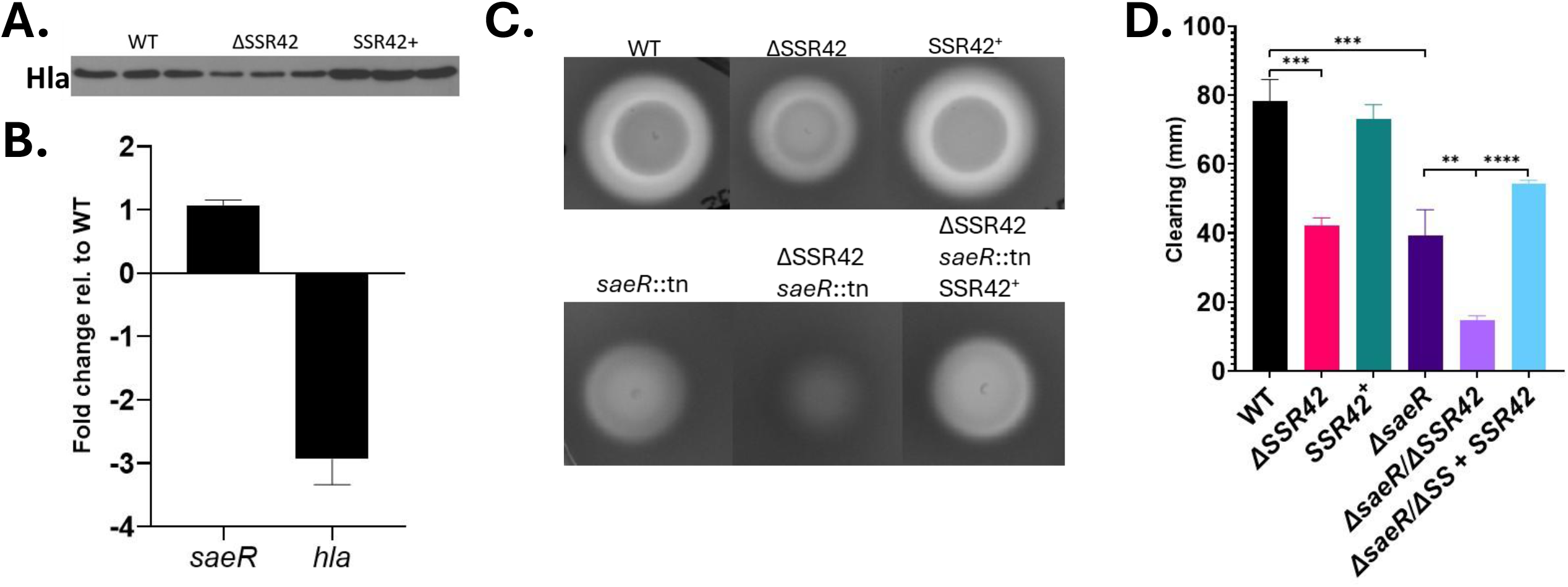
SSR42 regulates hemolysis independently of SaeRS. **(A)** Cultures were grown for 15h before secreted fractions were harvested, standardized to each other, and subjected to Western blot analysis to measure Hla abundance. **(B)** Wild-type and SSR42 mutant cultures were grown in triplicate for 15h before RNA was isolated and qPCR analysis was performed to assess for transcript abundance of *saeR* and *hla*. Error bars are ±SD. **(C)** Cultures were grown overnight and optical densities standardized before 7μl of culture was spot plated onto sheep blood agar and incubated at 37° for 15h. Image is representative of three biological replicates. **(D)** Quantification of C calculated using ImageJ software. All experiments were performed in biological triplicate. Error bars represent ±SEM. Statistical significance was assessed using Student’s t test and Welch’s correction (**; P < 0.01, ***; P < 0.001, ****P<0.0001).

### Complex and Opposing Regulation of *hla* transcription and translation by the Rsp-SSR42 Axis

To explore SSR42-mediated control of Hla, we first constructed a transcriptional reporter by fusing the *hla* promoter with luciferase and introduced this into our wild-type and SSR42 mutant strains. This revealed a decrease in *hla* promoter activity, especially during late stationary phase growth (the window of maximal SSR42 stability and impact) in the SSR42 mutant **(Figure 2A)**. Given that sRNAs typically manifest impact at the posttranscriptional level, we next explored if there was an indirect route to SSR42 control of *hla* expression. In our RNAseq dataset, expression of the transcription factor Rsp is increased (+4.3-fold) upon SSR42 deletion (29). Work by others in strain SH1000 demonstrates that Rsp directly activates *hla* expression, thus it’s upregulation upon deletion of SSR42 would not explain our findings (49). To confirm this, we inactivated *rsp* within our SSR42 mutant and monitored *hla* promoter activity.

**Figure 2.**
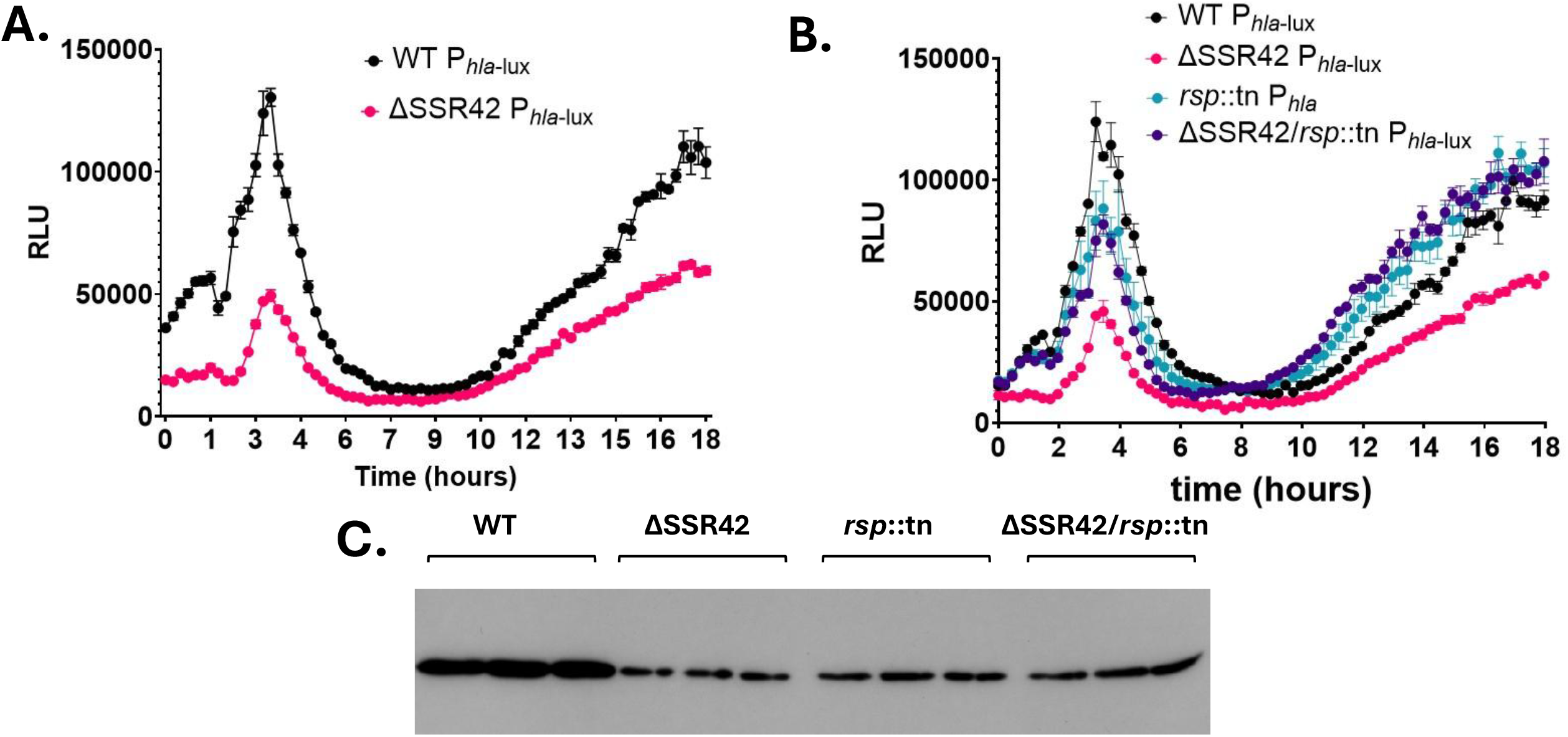
Complex transcriptional control of *hla* by the SSR42-Rsp axis. **(A)** P_*hla-lux*_ was expressed in the wild-type and SSR42 mutant strains with luminescence measurements taken every 15 min for 18h. **(B))** P_*hla-lux*_ was expressed in an *rsp* mutant as well as in the context of SSR42 deletion. Luminescence readings for A and B were measured using a Biotek Cytation5 plate reader. **(C)** Bacterial cultures were grown for 15h and supernatant samples were taken for Western blot analysis using antibodies specific to Hla. Blots were visualized using autoradiography. All bacterial cultures were grown in biological triplicate. Error bars are ±SEM.

Surprisingly, at odds with the work from SH1000, we found that deletion of *rsp* in an SSR42 mutant resulted in increased *hla* promoter activity that actually exceeds wild-type levels during stationary growth **(Figure 2B)**. To confirm if this transcriptional effect was carried over to the protein level, we next performed western blot analysis to assess abundance of Hla in the presence/absence of *rsp*. This revealed that instead of showing the expected increase in Hla abundance, the amount of Hla produced in an SSR42/*rsp* double mutantwas unchanged compared to the SSR42 single mutant **(Figure 2C)**. This suggests that the increased expression of *hla* in an SSR42/*rsp* mutant ultimately has no impact on Hla production as SSR42 is required for its effective translation. This is supported by our finding here that an *rsp* single mutant also has diminished Hla abundance, despite increased *hla* transcription, both as compared to WT. This is explained by the finding that SSR42 production is essentially ablated upon *rsp* deletion (50). Collectively, it appears that there is complex and opposing regulation of *hla*/Hla by the Rsp-SSR42 axis; with Rsp governing transcriptional control whilst SSR42 seemingly mediates translational activity.

### SSR42 posttranscriptionally controls Hla production through a discrete interaction site

To dissect the posttranscriptional control of Hla production by SSR42, we created a translational reporter by fusing the *hla* UTR and first few amino acids with GFP. This was then expressed under the control of a constitutive promoter to eliminate effects from transcriptional regulation. Here we found a significant decrease in the translation of Hla in the SSR42 mutant versus wild-type **(Figure 3A)**. Using a two plasmid system optimized to detect regulatory RNA control in gram-positive bacteria (51), we expressed SSR42 *in trans* alongside the constitutively transcribed *hla*::GFP fusion in the SSR42 mutant. In so doing, we note the restoration of Hla translation in the SSR42 mutant in comparison to an empty vector control strain **(Figure 3B)**. With these findings supporting a posttranscriptional, likely RNA-RNA binding event between SSR42 and *hla*, we next sought to narrow down the site of interaction between these two transcripts. To this end, we assessed the capacity of SSR42 to bind with the *hla* mRNA using the lntaRNA tool (52). In so doing, we identified a predicted RNA-RNA interaction between a region 291-nt downstream of the SSR42 +1 site and the *hla* 5’ UTR, 23-nt upstream from the RBS **(Figure 3C)**. To assess the specificity and validity of this prediction, we performed mutagenesis on the SSR42 binding site by both deleting and scrambling the DNA sequence and re-introducing this mutated complement into our SSR42 mutant for hemolysis assessment. This revealed that both mutated forms of the SSR42 complement resembled an SSR42 mutant strain, suggesting that the disrupted region is vital for SSR42’s regulation of hemolysis **(Figure 3D)**. To further validate the specificity of this binding region, we performed similar mutagenesis using our two-plasmid system. When our *hla*::GFP construct was expressed alongside native and mutated SSR42 we found that the scrambled allele was unable to restore Hla translation in comparison to the native form **(Figure 3E)**. Taken together, this corroborates interaction between the 5’ portion of SSR42 and the 5’ UTR of the *hla* mRNA and validates its importance for regulation of hemolysis through facilitation of translation.

**Figure 3.**
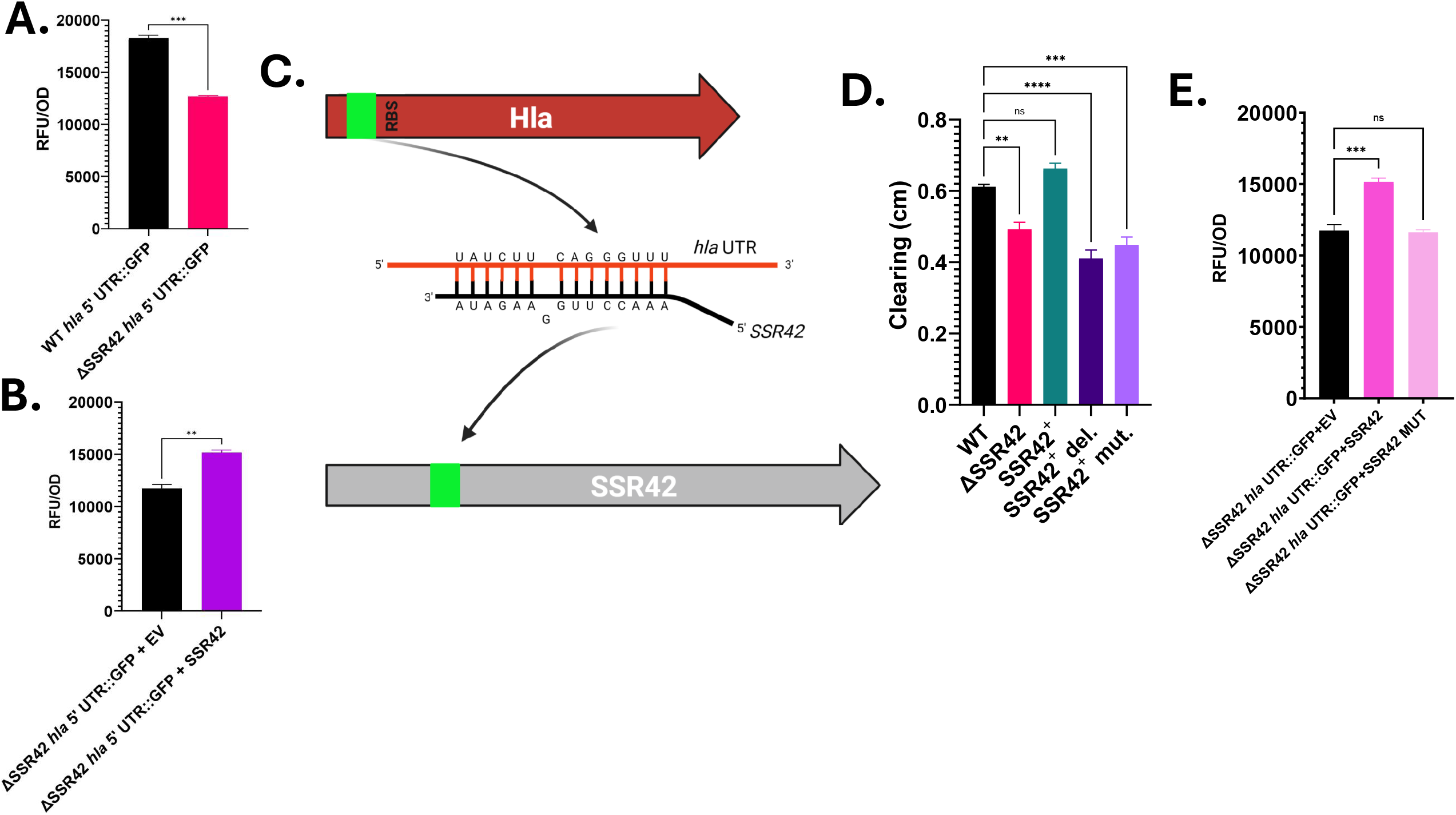
SSR42 posttranscriptionally controls Hla production through a discrete interaction site. **(A)** The *hla* 5’ UTR and first few amino acids were translationally fused to GFP and expressed via a constitutive promoter in the wild-type and SSR42 mutant. **(B)** As in A but the Hla reporter is expressed in an SSR42 mutant alongside either an EV (Empty Vector) or SSR42 *in trans*. **(C**) IntaRNA was used to identify thermodynamically likely regions of base paring between SSR42 and the *hla* mRNA. This identified an interaction region between the 5’ end of SSR42 and the 5’ UTR of *hla*. Created using Biorender.com. **(D)** The SSR42 complementation construct was mutagenized at the predicted site of interaction with *hla* and hemolysis was re-assessed. “Del.” refers to deletion of the predicted interaction site from the construct, while “mut.” corresponds to a scrambling of this sequence. Quantification of hemolytic activity was determined as in Figure 1CD. **(E)** The same predicted interaction site was scrambled in the two-plasmid system construct for SSR42 from B. The *hla 5’* UTR-GFP translational fusion was expressed alongside either an EV, WT SSR42, or scrambled SSR42 *in trans*. All experiments were performed in biological triplicate. Error bars are ±SEM. Statistical significance was assessed using Student’s t-test (ns, not significant, **; P < 0.01, ***; P < 0.001, ****; P < 0.0001).

### Stability of the *hla* mRNA is enhanced by the presence of SSR42

Regulatory RNAs typically wield positive impacts on target mRNAs by binding to and stabilizing them; resulting in increased capacity for translation. To determine if something similar occurs between SSR42 and the *hla* mRNA, we performed transcriptional arrest on the wild-type, SSR42 mutant and complementing strains and quantified *hla* mRNA abundance over time following treatment with rifampin. In so doing we determined that the *hla* mRNA exhibited significant de stabilization in the SSR42 mutant, which was profoundly reversed upon reintroduction of SSR42 **(Figure 4)**. Specifically, the *hla* mRNA half-life is 7.63 minutes in the wild-type whereas in an SSR42 mutant this is decreased to 5.46 minutes. Taken together, this data suggests that SSR42 plays a role in stabilizing the *hla* mRNA resulting in enhanced downstream translation of active alpha-toxin.

**Figure 4.**
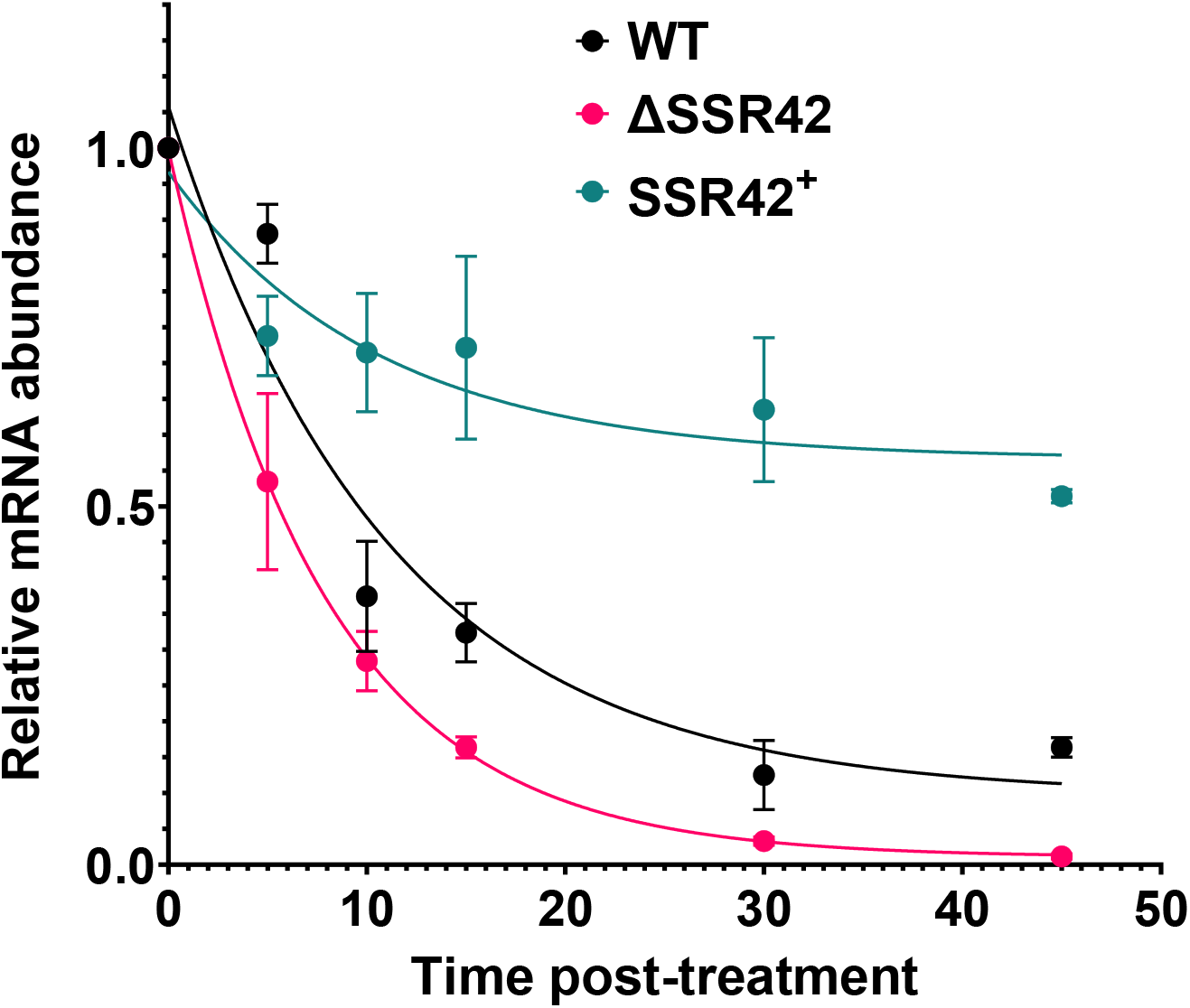
SSR42 is responsible for stabilizing the *hla* mRNA. Bacterial cultures were grown for 15h before the addition of Rifampin. Samples were withdrawn at the indicated timepoints and RNA isolated for downstream qPCR to quantify RNA abundance and relative mRNA stability in comparison to T=0. All RNA quantifications were normalized to 16S rRNA and a one phase decay curve was generated using Graphpad Prism. All bacterial cultures were grown in biological triplicate. Error Bars represent

### SSR42 binds the *hla* 5’ UTR directly but likely requires additional facilitating factors

To fully validate post-transcriptional regulation of *hla* by SSR42, we performed RNA EMSAs to assess binding. First, we *in vitro* transcribed and radiolabeled the *hla* 5’ UTR fragment and incubated it with increasing concentrations of full-length SSR42. In so doing we observed a modest but detectable, concentration-dependent shift in size indicating these two RNA molecules interact **(Figure 5)**. When the same experiment was repeated using a fragment of *splE*, which is known not to interact with SSR42 (29), we observed no such interaction. Importantly, when we incubated *hla* mRNA with SSR42 containing the mutagenized or deleted sequences from **Figure 3**, the observed shift completely disappears, corroborating the specificity of this binding event. As such, it is clear that SSR42 binds discretely and specifically to native SSR42, however there may well be additional factors required for robust interaction and binding between these molecules.

**Figure 5.**
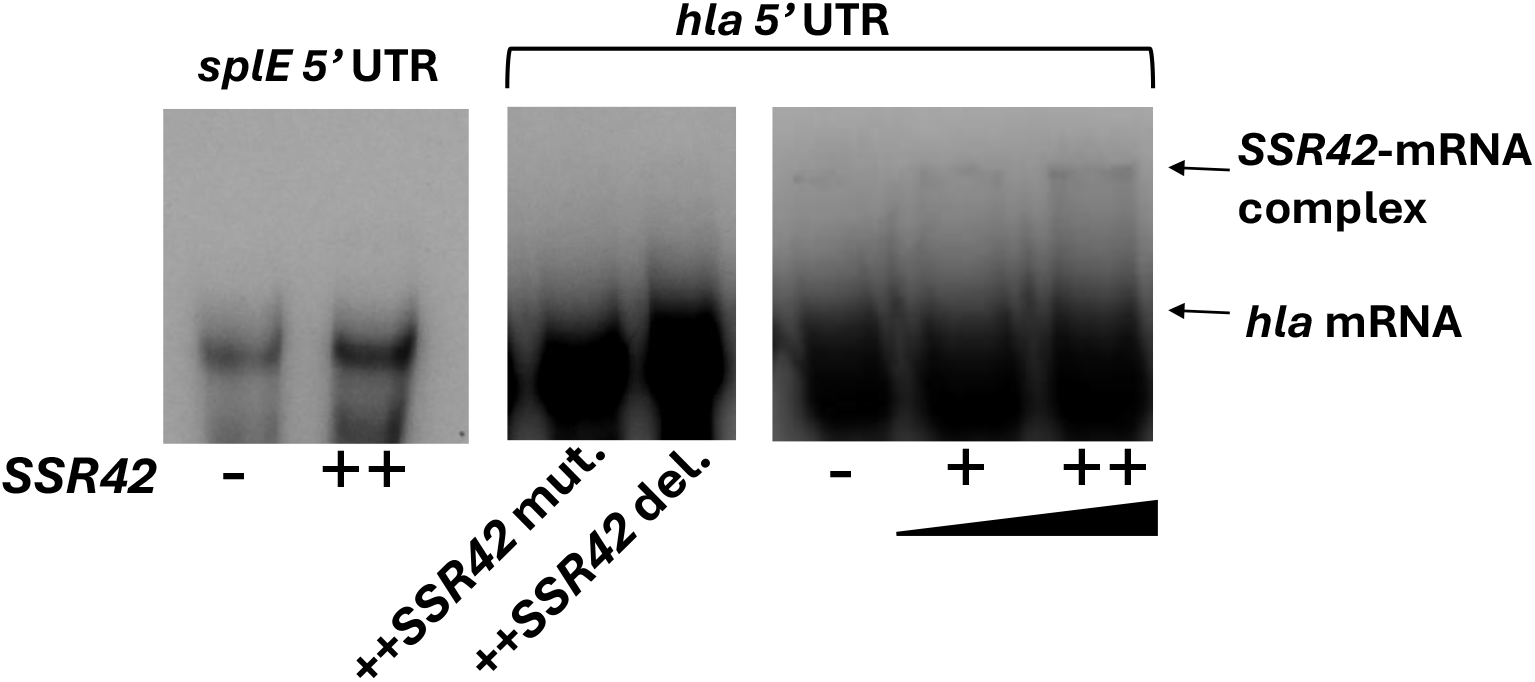
SSR42 binds the *hla* mRNA directly but may require additional factors. *in vitro* transcribed SSR42 was added in increasing amounts (+=200-fold excess, ++=500-fold excess) to radiolabeled portions of the *hla* mRNA. Samples were heated to 90°C for 2 min and then incubated at room temperature for 30 min before being loaded on to a non-denaturing gel. “Del.” and “mut.” are as in Figure 3. At identical concentrations, no binding is seen for radiolabeled *splE*. Image is representative of 3 independent replicates.

## Discussion

In this study, we investigate the relationship between SSR42 and the key hemolysin Hla. The link between these two factors has been well established in previous studies (35, 36), however, the regulatory route to control has yet to be fully elucidated. Work by Horn et al., in strain 6850 suggested that SSR42 exerts its effects on *hla*/Hla production in an indirect manner, acting through the virulence controlling two-component system SaeRS (35). This is clearly contrary to our findings herein. The disparity between SSR42’s role in hemolysis regulation in the strain 6850 and our own USA300-based studies may suggest SSR42 exhibits strain-specific behavior. With that said, SSR42’s role as an activator of hemolysis is consistent across diverse lineages, including UAMS-1 (ST-30, MSSA), 2012-3 (ST398, MRSA), 6850 (ST50, MSSA), and in USA300 strains LAC and HOU (ST-8, MRSA) (29, 34-36). Moreover, work by the Dunman group found (as we do) no alteration in *saeRS* expression in SSR42 mutants of strain LAC or UAMS-1 (36). Others have noted that the lineage 6850 belongs to (ST50) is as an apparent singleton, with little similarity to common clonal complexes (54). Therefore, it is perhaps unsurprising that the SSR42 regulon in this phylogenetically divergent lineage may be different from that which is observed in more commonly found sequence types.

A consideration with our findings is that although we show a clear impact of SSR42 on *hla* mRNA stability and alpha-toxin protein production, our molecular interaction studies using RNA-RNA EMSAs only result in a modest shift. This suggests that the *in vitro* assessment used herein may be missing factor(s) that are required for complete and robust binding of these two molecules. For example, despite efforts to control and stabilize the generated RNA molecules, it is still likely that the secondary structure within our *in vitro* conditions will be altered in comparison to that seen *in vivo*. As secondary structure inevitably impacts binding events, this could be one caveat that prevents our EMSA results from reaching their fullest significance. Moreover, additional factors may be required for full binding that are only present within the bacterial cell. RNA chaperones are the biggest possibility here; with Hfq being a prime example that has been widely studied across species (55). However, the role for Hfq in *S. aureus* remains nebulous - some studies identified a binding event between RNAIII and Hfq, but this interaction did not lead to enhanced complex formation between RNAIII and its target mRNAs. (56, 57) Moreover, Hfq is very lowly expressed in *S. aureus* in comparison to other bacterial species, thus, while its role is not completely understood, it is unlikely to govern SSR42 activity. There are no known Hfq homologs in *S. aureus*, although there is some work in *B. subtilis* that suggests additional RNA chaperones exist that are as yet uncharacterized (56). Thus, the possibility of an RNA chaperone facilitating SSR42-*hla* interactions cannot be eliminated without further study. Another missing component aiding this interaction could come from sRNA-sRNA interactions, in which regulatory RNAs bind to each other to either facilitate or repress regulatory behavior. One study utilizing the RNaselll CLASH technique to discover novel RNA chimeras in *S. aureus* reported that 7% of all RNA interactions were between sRNAs and other sRNAs (58) Although some of these interactions have been reported to be RNA ‘sponging’ events, as is the case with RsaE and Rsal for example, the potential for sRNA-sRNA interactions is understudied and could possibly be playing a role herein.

Thus, although *in vitro* systems have strong utility, they can be limited in their ability to capture delicate RNA interactions. As such, sRNA studies are perhaps best performed using optimized translational reporter systems within bacterial cells, such as those used herein to study SSR42-*hla* engagement (59).

In addition to controlling *hla* mRNA stability, we also show that SSR42 exerts influence over P_*hla*_, seemingly through its impact on Rsp production. Importantly, alongside Rsp, there are 2 other transcriptional regulators with altered expression in the SSR42 mutant (29): SarZ (+1.98-fold) and SarR (-7.06-fold). To ensure these factors do not present additional indirect nodes by which SSR42 can control *hla*, we next considered their influence. In the context of SarZ, it has been shown to activate Agr (60), thus, it’s increased abundance in an SSR42 mutant should result in increased Agr activity (not observed), which would in turn upregulate *hla;* thus, this route would not appear to be causative. Indeed, deleting *sarZ* in an SSR42 mutant does not result in changes to *hla* promoter activity **(Figure S2A)**, confirming this contention. Regarding SarR, in strain SH1000 it outcompetes SarA for binding to P2 of the agroperon, subsequently repressing Agrfunction (47, 61). Thus, it’s significant downregulation in an SSR42 mutant would upregulate Agr activity, and thus *hla* expression; which again we do not see. To confirm this, we counteracted the downregulation of *sarR* in the SSR42 mutant by overexpressing it and did indeed observe an additional decrease in *hla* expression during stationary phase growth **(Figure S2B)**. Thus, it would appear that the only transcriptional effect SSR42 has on P_*hla*_ is wielded via Rsp, however the direct action of SSR42 on *hla* mRNA stability is dominant to any alterations in expression observed.

An intriguing facet of SSR42-mediated regulation of hemolysis is its similarity to the well-characterized mechanism by which RNAIII interacts with the *hla* mRNA(20). Indeed, both regulators are long (RNAIII ∼500-nt, SSR42 ∼1,200-nt) stable RNA molecules that bind the *hla* transcript upstream of the RBS and activate translation. RNAIII binds the *hla* mRNA and alters its secondary structure, allowing for ribosomal access to the RBS (20). Importantly, it has not been shown that RNAIII binding affects stability of the *hla* mRNA, however. Herein, we show that this is exactly the effect SSR42 has - it’s binding to the *hla* mRNA increases stability and seeming capacity for Hla translation. As these two factors bind within the same region of the 5’ UTR of *hla*, (shown in **Figure S3)**, we suggest a fascinating scenario whereby two large regulatory RNA molecules may be required to ensure the production of a vital virulence factor through different but complementary mechanisms. Interestingly, RNAIII has been shown in the literature to be part of a similar dual-action partnership, where it acts alongside the regulatory RNA SprD to ensure the production of the immune evasion protein Sbi (63) Specifically, RNAIII and SprD are produced at different intervals temporally, which ensures Sbi production during transitional phases of growth. Examining RNAIII and SSR42’s expression reveals the potential for a similarly timed regulatory program controlling Hla. RNAIII reaches peak expression once quorum is achieved, typically during the switch to stationary phase (63, 64), while SSR42 is most abundant in deeper stationary phase (36). Therefore, it is possible that RNAIII’s earlier activation of *hla* is during the switch by *S. aureus to* toxin production, while SSR42 continues to stabilize the transcript during prolonged growth to ensure Hla production. Moreover, this handoff could also be based on environmental conditions, as SSR42 is induced by host-like conditions (29), which may be facilitate continued Hla production during infection.

Although the SSR42 transcript has previously been shown to be 98% conserved at the sequence level, it has been annotated under many different names and has even been reported to be expressed in different isoforms depending on the strain (34, 36). Indeed, previous work in an ST398 isolate of *S. aureus* revealed a short isoform of SSR42 (therein called RsaX28) that corresponds to ∼500nt fragment within the middle of the ∼1,200nt SSR42 transcript found in USA300 strains. Interestingly, that work identified a relationship between this molecule and RNAIII in which a binding event stabilized the Agr effector, leading to downstream activation of hemolysis (34). It is thus pertinent to determine if this event is impacting *hla* production in the USA300 background, or if this is once again a strain-specific and/or SSR42-isoform specific behavior. First and most importantly, neither our previously work nor published SSR42 microarray studies have identified any changes to the RNAIII transcript or Hid protein abundance upon deletion of SSR42 (29,36). Nevertheless, we quantified RNAIII half-lives in our strains following transcriptional arrest to be sure there is no influence of SSR42 on Agr activity. In so doing we identified no changes in RNAIII transcript stability in the absence of SSR42 that would suggest that this same binding event is occurring **(Figure S4)**. Thus, we can conclude that the behavior displayed by the highly unique isoform of SSR42 identified in the ST398 strain is not occurring in USA300 strains and is not playing a role in SSR42-mediated regulation of hemolysis.

We propose that SSR42 is a novel, RNA-based direct regulator of alpha-toxin activity that acts independently of canonical pathways such as SaeRS or Agr. To our knowledge, this is the only other regulatory RNA that has been identified to directly impact *hla* other than RNAIII. We posit that these two large regulatory molecules maintain independent binding interactions with *hla* but may work in concert with each other to ensure the production of this key virulence factor. These results build on earlier studies of SSR42 and highlight its role in coordinating multiple regulatory inputs that govern virulence factor expression **(Figure 6)**. Taken together, our findings support the continued emergence of SSR42 as a highly impactful regulator of *S. aureus* virulence, and sheds light into still understudied facets of the regulation of pathogenesis, especially as it pertains to RNA-based regulation.

**Figure 6.**
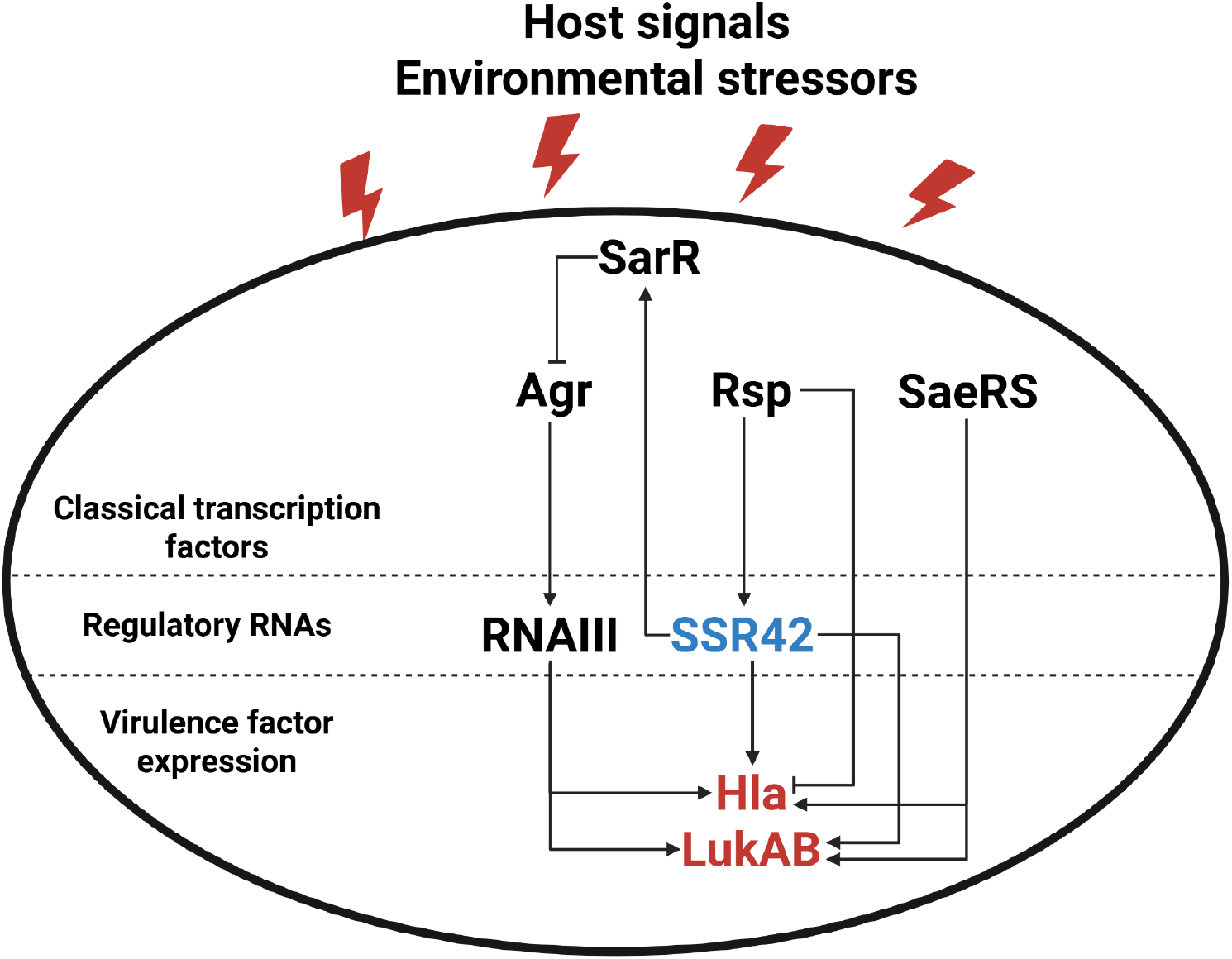
Multi-tier regulation of virulence factor expression in *Staphylococcus aureus*. In response to environmental stimuli, transcriptional regulators directly impact virulence factor expression but also control the activity of central regulatory RNAs SSR42 and RNAIII. The overlapping, yet independent pathways of these regulatory transcripts allow for tightly modulated control over protein expression and ensured activation of key virulence determinants. Previously characterized regulatory pathways are shown with SSR42 highlighted in blue as a central focus of this study and virulence factors of interest highlighted in red.

## Acknowledgements

This study was supported by grant AI188698 (L.N.S.) from the National Institute of Allergy and Infectious Diseases.

## Notes

### Competing Interest Statement

The authors have declared no competing interest.

