## Supplemental Figures and Tables for "SSR42 Enhances the Hemolytic Capacity of *Staphylococcus aureus* by Stabilizing the *hla* mRNA"

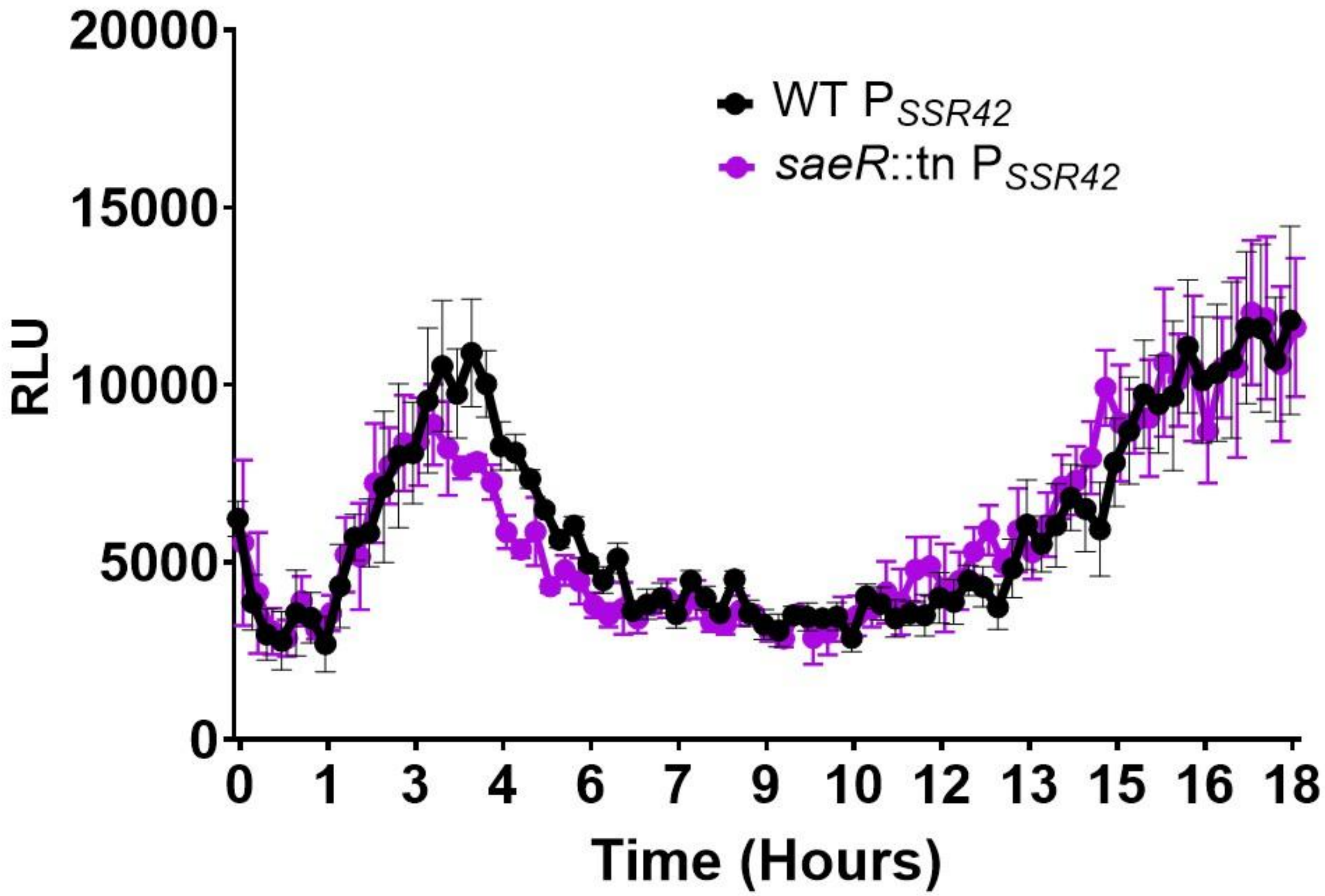


**Figure S1. SaeRS does not control $P_{SSR42}$.** The SSR42 promoter was fused to luciferase and expressed in the wild-type and *saeR* mutant strains, with fluorescent measurements taken every 15 min for 18h. Bacterial cultures were grown in biological triplicate, with error bars shown ±SEM.

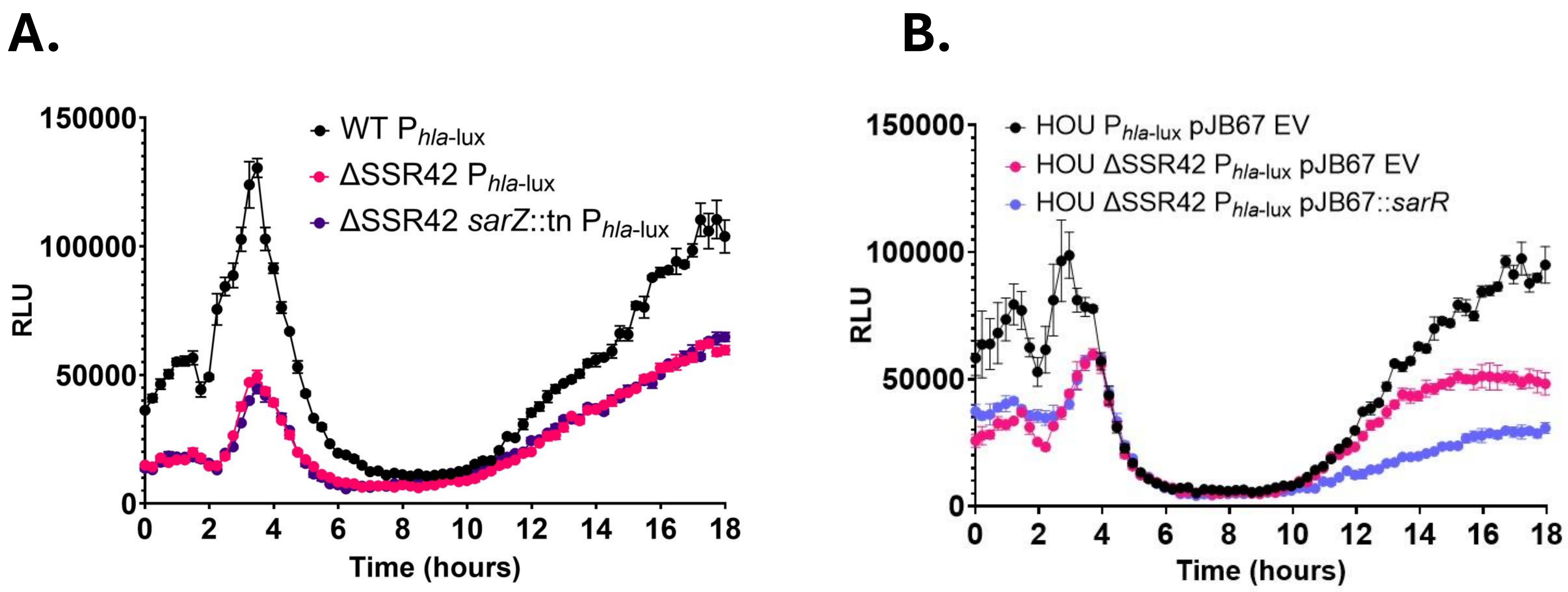


**Figure S2 . Alterations in SarZ and SarR abundance in the SSR42 mutant are not causative for decreased *hla* transcription. (A)** $P_{hla\text{-lux}}$ was expressed in a *sarZ* mutant in the context of SSR42 deletion. Luminescence was measured every 15 min for 18h. **(B)** As in A but utilizing a *sarR* overexpression strain in the context of SSR42 deletion. Bacterial strains were grown in biological triplicate. Error Bars are ±SEM

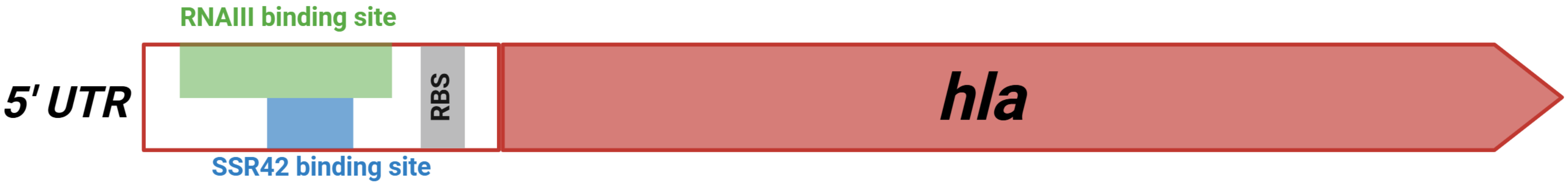


**Figure S3. Overlapping regulation of *hla* by regulatory RNA molecules in *S. aureus*.** Binding sites for RNAIII and SSR42 are represented alongside other features of interest on the *hla* mRNA.

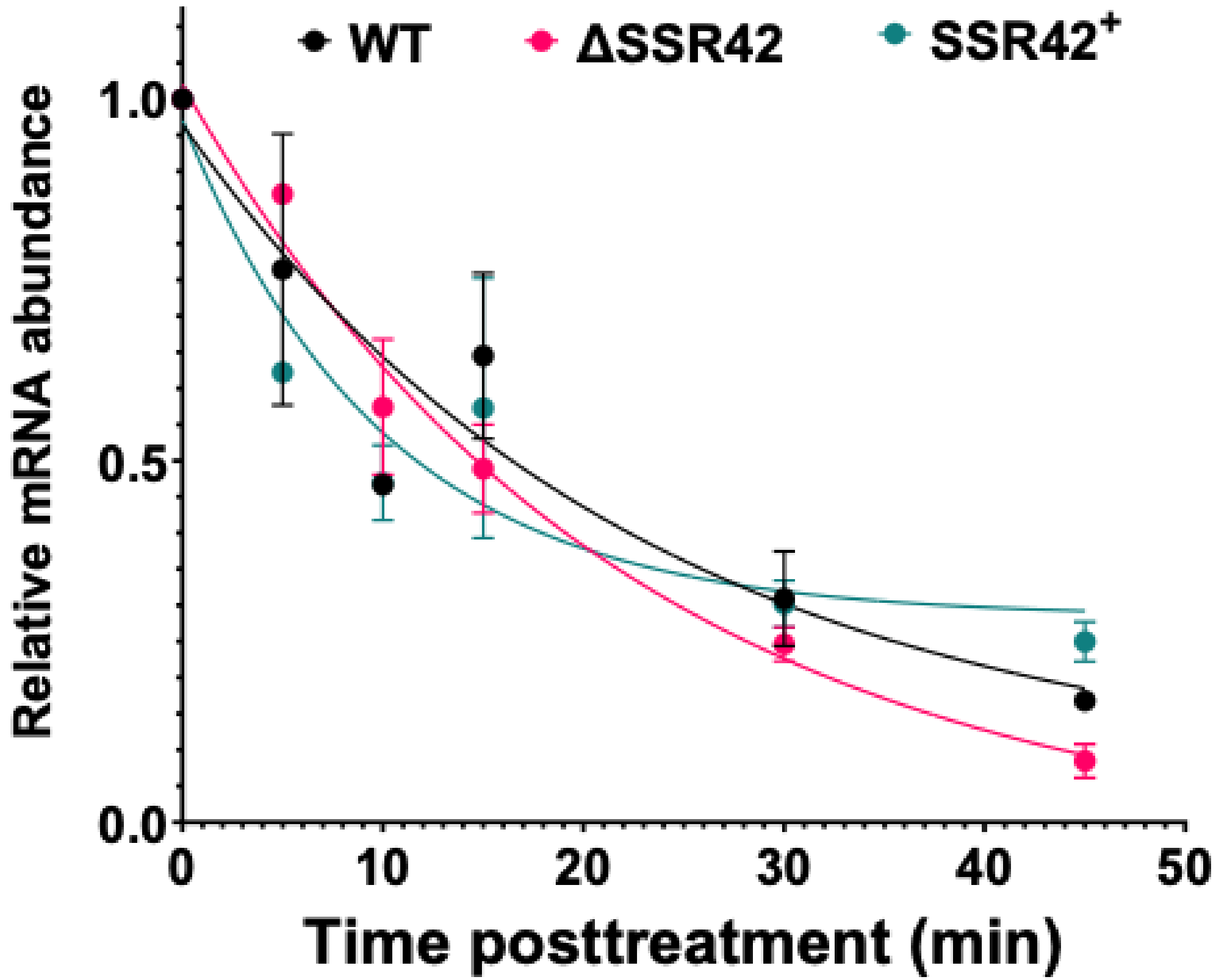


**Figure S4. RNAIII is not impacted by mutation of SSR42.** Bacterial cultures were grown for 15h before Rifampin was added and samples were extracted at the time points shown. RNA was then harvested, and qPCR was used to assess stability of RNAIII. RNA abundance was normalized to 16S rRNA and a one phase decay curve was generated using Graphpad Prism. All bacterial cultures were grown in biological triplicate. Error Bars represent ±SEM.

**Table S1: Bacterial strains used in this study.**

| Strain Name | Description | Source |
|---|---|---|
| ***E. coli*** | | |
| DH5α | Cloning strain | (1) |
| ***S. aureus*** | | |
| RN4220 | Cloning strain, restriction-deficient | Lab stock |
| TCH1516 | USA300 HOU CA-MRSA Erm$^s$, Wildtype parent strain | (2) |
| BRT2544 | HOU ΔSSR42 | (3) |
| LNS1927 | HOU pMK4 EV | (3) |
| BRT2545 | HOU ΔSSR42 pMK4 EV | (3) |
| BRT2546 | HOU ΔSSR42 pMK4::SSR42 | (3) |
| BDG2622 | HOU *saeR*::tn | This study |
| BRT3016 | HOU ΔSSR42 *saeR*::tn | This study |
| BRT3039 | HOU saeR::tn pMK4 EV | This study |
| BRT3040 | HOU ΔSSR42 *saeR*::tn pMK4 EV | This study |
| BRT3017 | HOU ΔSSR42 *saeR*::tn pMK4::SSR42 | This study |
| MEJ3545 | HOU pXen-1::$P_{hla}$ | This study |
| MEJ3549 | HOU ΔSSR42 pXen-1::$P_{hla}$ | This study |
| MEJ4048 | HOU pCN33::*hla* UTR::GFP | This study |
| MEJ4049 | HOU ΔSSR42 pCN33::*hla* UTR::GFP | This study |
| MEJ4050 | HOU ΔSSR42 pCN33::*hla* UTR::GFP, pICS3 EV | This study |
| MEJ4051 | HOU ΔSSR42 pCN33::*hla* UTR::GFP, pICS3::SSR42 | This study |
| MEJ4052 | HOU *sarZ*::tn pXen-1::$P_{hla}$ | This study |
| MEJ4053 | HOU ΔSSR42 sarZ::tn pXen-1::$P_{hla}$ | This study |
| MEJ4054 | HOU Δ*sarR* pXen-1::$P_{hla}$ | This study |
| MEJ4055 | HOU Δ*sarR* ΔSSR42 sarZ::tn pXen-1::$P_{hla}$ | This study |
| MEJ4141 | HOU pXen-1::$P_{hla}$ pJB67 EV | This study |
| MEJ4464 | HOU ΔSSR42 pXen-1::$P_{hla}$ pJB67 EV | This study |
| MEJ4056 | HOU pXen-1::$P_{hla}$ pJB67::*sarR* | This study |
| MEJ4057 | HOU ΔSSR42 pXen-1::$P_{hla}$ pJB67::*sarR* | This study |

| | | |
|---|---|---|
| MEJ4058 | HOU *rsp*::tn pXen-1::P*hla* | This study |
| MEJ4059 | HOU ΔSSR42 *rsp*::tn pXen-1::P*hla* | This study |
| MEJ4598 | HOU pXen-1::*PSSR42* | This study |
| MEJ4599 | HOU *saeR*::ten pXen-1::*PSSR42* | This study |
| MEJ4060 | HOU ΔSSR42 pMK4::SSR42 MUT | This study |
| MEJ4061 | HOU ΔSSR42 pMK4::SSR42 DEL | This study |
| MEJ4062 | HOU ΔSSR42 pCN33::*hla* UTR::GFP, pICS3::SSR42 MUT | This study |

**Table S2: Plasmids used in this study.**

| Plasmids | Description | Source |
|---|---|---|
| pXen-1 | Luciferase reporter plasmid, *E. coli* $Amp^R$, *S. aureus* $CM^R$ | (4) |
| pCN33 | mRNA target GFP reporter, *E. coli* $Amp^R$, *S. aureus* $Ery^R$ | (5) |
| pICS3 | RNA expression vector, *E. coli* $Amp^R$, *S. aureus* $CM^R$ | (5) |
| pJB67 | pCN51 with optimized RBS and Cd-inducible promoter, *E. coli* $Amp^R$, *S. aureus* $Ery^R$ | (6) |

The following abbreviations refer to antibiotic resistance cassettes: Ery, erythromycin; Tet, tetracycline; CM, chloramphenicol; Amp, ampicillin.

**Table S3: Primers used in this study.**

| Primer | Sequence | Description |
|---|---|---|
| OL2888 | ACT<u>GGATCC</u>GTGTCTATAATACTTTGACCT | SSR42 complementation F |
| OL3734 | ATG<u>GTCGAC</u>GTTCAATATTTACTACAAAGTCGTG | SSR42 complementation R |
| OL3114 | CACCTCTGTTCTTACGACCTC | *saeR* screening F |
| OL3115 | GGTGTTATCAATTAGAAGTCAAATC | *saeR* screening R |
| OL2393 | TCGTATGTTGTGTGGAATTG | pMK4 MCS F |
| OL2394 | GTGCTGCAAGGCGATTAAG | pMK4 MCS R |
| OL398 | TCCTACGGGAGGCAGCAG | 16S F |
| OL399 | GGACTACCAGGGTATCTAATCCTGTT | 16S R |
| OL3116 | ACCACAATAACTCAAATTCCTTAATACG | *saeR* RT-qPCR F |
| OL3117 | GTTGAACAACTGTCGTTTGATGA | *saeR* RT-qPCR R |
| OL4036 | GGCTCTATGAAAGCAGCAGATA | *hla* RT-qPCR F |
| OL4037 | CTGTAGCGAAGTCTGGTGAAA | *hla* RT-qPCR R |

| | | |
|---|---|---|
| OL843 | ATG<u>GAATTC</u>CCCGACGAAATTCCAAAC | *hla* promoter for pXen-1 F |
| OL844 | ATG<u>GGATCC</u>AAGGCCAGGCTAAACCAC | *hla* promoter for pXen-1 R |
| OL4526 | ATG<u>GAATTC</u>GTTGGCATGTCATATTTCTCTCTCCTG | SSR42 promoter for pXen-1 F |
| OL7967 | AATT<u>GAATTC</u>CATTGCGAACAATATAAAAGTAAAATTAGAAAGA | SSR42 promoter for pXen-1 R |
| OL5416 | ATCAGAGCAGATTGTACTGAG | pXen-1 MCS F |
| OL5417 | ACTCCTCAGAGATGCGAC | pXen-1 MCS R |
| OL7424 | AATT<u>AGATCT</u>TTCCCGACGAAATTCCAAAC | *hla* UTR for pCN33 F |
| OL7481 | AATT<u>GATATC</u>GGAACCTAGCAATAGTGTTGTTG | *hla* UTR for pCN33 R |
| OL7322 | GGGAGAAAGGCGGACAG | pCN33 MCS F |
| OL7323 | GCACGCGTCTTGTAAGTC | pCN33 MCS R |
| OL5365 | CGATGAAATCAGTACTTGACAAC | *sarZ* screening F |
| OL5366 | ACATATCGATGCATACTTCTGC | *sarZ* screening R |
| OL4577 | GACTAGTGTACCTTGTTTCAAGC | *sarR* screening F |
| OL4578 | TTGCTACAACAAGATGTGCATC | *sarR* screening R |
| OL6053 | GTTCGCAATGCATAAAAACAAGCG | *rsp* screening F |
| OL6054 | GGTGTACTGATTAAATCGGCAATTAATGAC | *rsp* screening R |
| OL6637 | AATT<u>CATATG</u>AGTAAAATTAATGACATTAATG | *sarR* for pJB67 F |
| OL6640 | AATT<u>GAATTC</u>CCCATCACCTTATGATAGGGAG | *sarR* for pJB67 R |
| OL4093 | GGCAGATAATGATGATCGC | pJB67 MCS F |
| OL4094 | CATGTCAACGATAATACAAAATATAATAC | pJB67 MCS R |
| OL8046 | **acca**TTGATTTGAATAATTTGTAGCG | SSR42 mut. F |
| OL8047 | **ccgcg**CCATCAACTAAGAAATTTTAAACAATC | SSR42 mut. R |
| OL8044 | CCATCAACTAAGAAATTTTAAACAATC | SSR42 del. F |
| OL8045 | TTGATTTGAATAATTTGTAGCG | SSR42 del. R |
| OL7499 | AATT*TAATACGACTCACTATAGGG*GATTTCAAACCTATGTATTTC | SSR42 + T7 promoter EMSA F |
| OL7500 | AATATGATGTTCAATATTTACTACAAAGTCG | SSR42 EMSA R |
| OL7523 | AATTT*AATACGACTCACTATAGGG*GCAGATAATTTAGATAAATAAATCATCCATCCATA | *splE* 5’ UTR + T7 EMSA F |
| OL7524 | TACCCTCAACCACTGTTGT | *splE* 5’ UTR EMSA R |
| OL8118 | AATT*TAATACGACTCACTATAGGG*CTCATCATCACTCAGTAATTTATCAGTTG | *hla* 5’ UTR + T7 promoter EMSA F |
| OL8117 | CAGAATCTGCGGCATTAGC | *hla* 5’ UTR EMSA R |

*underline, restriction enzyme site; bold, mutagenesis region; italics, T7 promoter.
